# Seeing and Extrapolating motion trajectories share common informative activation patterns in primary visual cortex

**DOI:** 10.1101/2022.05.26.493554

**Authors:** Camila Silveira Agostino, Christian Merkel, Felix Ball, Peter Vavra, Hermann Hinrichs, Toemme Noesselt

**Affiliations:** Department of Biological Psychology, Otto-von-Guericke-University Magdeburg, Magdeburg, Germany; European Structural and Investment Funds-International Graduate School (ESF-GS) Analysis, Imaging, and Modelling of Neuronal and Inflammatory Processes (ABINEP) International Graduate School, Otto-Von-Guericke-University, Magdeburg, Germany; Department of Neurology, Otto-von-Guericke University, Magdeburg, Germany; Center for Behavioral Brain Sciences, Otto-von-Guericke-University Magdeburg, Germany; Department of Behavioral Neurology, Leibniz Institute for Neurobiology, Magdeburg

**Author notes:** **Corresponding Author:** Camila S. Agostino –. **Ethics statement** This study was approved by the ethics committee of the Otto-von-Guericke-University. **Data availability statement** Data will be shared upon request with the need for approval from the requesting researcher’s local ethics committee.

**Keywords:** Dynamic occlusion, V1, fMRI, MVPA, Retinotopic Mapping

## Abstract

The natural environment is dynamic and moving objects become constantly occluded, engaging the brain in a challenging completion process to estimate where and when the object might reappear. Although motion extrapolation is critical in daily life – imagine crossing the street while an approaching car is occluded by a larger standing vehicle – its neural underpinnings are still not well understood. While the engagement of low-level visual cortex during dynamic occlusion has been postulated, most of the previous group-level fMRI-studies failed to find evidence for an involvement of low-level visual areas during occlusion. In this fMRI-study, we therefore used individually-defined retinotopic maps and multivariate pattern analysis to characterize the neural basis of visible and occluded motion in humans. To this end, participants learned velocity-direction pairings (slow motion-upwards; fast motion-downwards or vice versa) during a training phase without occlusion and judged the stimulus direction, based on its velocity, during a following test phase with occlusion. We find that occluded motion direction can be predicted from the activity patterns during visible motion within low-level visual areas, supporting the notion of a mental representation of motion trajectory in these regions during occlusion.

**Highlights:**

* Dynamically occluded information is processed in low-level visual cortex
* Specific regions inside low-level visual areas encode visible and dynamically occluded information
* Overlap of visible and occluded informative activity patterns in the visual field suggest shared computational circuits in primary visual cortex

## 1. INTRODUCTION

In our daily life, we often miss critical input from our visual environment: simple eye blinks, occlusion of moving objects and other internal and external disruptive changes may fragment part of the incoming information. The human brain has evolved to make adequate inferences about these missing inputs (for review, see e,g, Thielen et al., 2019). For instance, when driving or crossing a street, we are able to estimate the time when a vehicle will reappear, after being occluded by a bus, to plan our next action accordingly, indicating that we can successfully infer the reappearance of a dynamically occluded object (e.g. Coull et al, 2008, Dittrich & Noesselt, 2018). Although such inference mechanisms underpin many actions of our daily life, little is known about their exact neural representations and their informational content. In this study, we used brain imaging to identify the low-level visual regions instrumental in the processing of dynamically occluded objects. In particular, we use functional magnetic resonance imagining (fMRI) and multivariate pattern analysis (MVPA) to compare the encoding of occluded and visible information in individually-defined low-level visual regions.

Engagement of low-level visual areas in processes related to motion prediction remain ambiguous. For instance, a recent fMRI study (Ekman, Kok & de Lange, 2017) on *apparent* motion reported that enhanced fMRI-responses in V1 could be triggered by the sole presentation of the first stimulus of a series of spatially-distinct flashes. Remarkably, V1 activity resembled the entire stimulus sequence even though the subsequent visual input was not present; the activity pattern during prediction was temporally compressed, suggesting that V1 anticipates the presence of the expected targets. In the realm of *continuous* motion extrapolation, behavioural studies also suggest the engagement of V1 during dynamic occlusion of a moving object; and several hypotheses have been proposed to account for the mechanisms underlying motion prediction. One of the most common hypotheses postulates that time-to-contact (TTC, DeLucia & Liddell, 1998) estimation of the occluded object would engage early stages of visual processing by using a mental representation of the visual trajectory (de’Sperati & Deubel, 2006; Battaglini et al, 2014) and memory of temporal information acquired during watching the visible trajectory (Khoei, Masson, & Perrinet, 2013; Makin et al., 2008; Makin & Bertamini, 2014; Makin & Poliakoff, 2011; Makin et al., 2012; Battaglini et al., 2013). Additionally, enhanced attentional resource allocation was already observed at locations that contain temporarily occluded moving targets in behavioural studies (Scholl, 1999; Flombaum, Scholl, Pylyshyn, 2008); in accord, an anisotropic distribution of representational enhancement was found in the direction of predicted motion (Verghese & McKee, 2002; Atsma, Koning & van Lier, 2012; Frielink-Loing, Koning, van Lier, 2017). Importantly, such attentional extrapolation can usually only be observed with a very low number of simultaneously relevant trajectories (Keane, Pylyshyn, 2006; Zhong, et al, 2014; Vul, et al, 2009). These behavioural studies all show modulations along a spatial gradient relative to the location of the extrapolated object, thus pointing at an involvement of retinotopically organized areas, i.e. low-level visual cortex, in motion extrapolation.

In contrast, previous fMRI-studies focusing on the neural underpinnings of occluded moving objects often failed to observe evidence for the involvement of low-level visual areas and rather observed a recruitment of parietal regions especially intraparietal sulcus (IPS), (Shuwairi et al., 2007; O’Reilly, Mesulam, & Nobre, 2008); or reported decreased fMRI-signals in these regions instead (Olson et al., 2003). An alternative hypothesis, in accord with this reduction of fMRI-signal in low-level visual areas, would be that inference of predictable trajectories reduce neural activity, similar to signal decrease in other highly predictable environment in a variety of tasks (Alink et al, 2010, Krala, et al., 2019, van Heusden et al., 2019). This hypothesis is in line with Rao and Ballard’s (1999) hierarchical predictive coding model, which postulates that feedback and feedforward connections convey predictions to lower levels and error estimates to higher levels, respectively, and that deviations from a predicted outcome would lead to enhanced signalling in low-level areas.

Another reason why many of previous fMRI-studies on the neural basis of motion extrapolation may have failed to observe the involvement of low-level visual areas may be grounded in conceptual and methodological issues. The exact anatomical location of functionally distinct visual areas is highly variable across humans (Greenlee, 2000; Amunts, 2000), hence, any effects may be diminished when using standard voxel-based group mean analyses, as it was done by most previous investigations (Olson et al., 2003; Shuwairi et al., 2007; O’Reilly, Mesulam, & Nobre, 2008). So far, there are few studies which investigated dynamic occlusion used retinotopic maps to identify subject-specific regions of interest and observed modulations of fMRI signal in low-level visual areas. Recently, Erlikhman and Caplovitz (2017) used subject-specific retinotopic mapping to identify subject-specific primary visual cortex together with multivariate pattern analysis (MVPA) to test whether differences in the shape of dynamically occluded objects moving along a single trajectory are already decodable in low-level visual areas. The authors reported enhanced activation in V1 during occlusion, but failed to observe evidence for the objects’ shape in the activity patterns in this region.

To our knowledge, no study so far investigated whether the neural representation of different extrapolated trajectories can already be decoded in low-level visual areas. Therefore, the main aim of this study was to differentiate patterns of activity in individually defined low-level visual cortex for different visible and occluded trajectories during the presentation of dynamically occluded stimulus. To this end, we employed a prediction motion paradigm (Hecht & Savelsbergh, 2004, Battaglini & Ghiani, 2021), where participants learned specific motion velocity-trajectory associations during an initial familiarisation phase based on visible motion and had to judge time- and point-of-contact during a test phase in which the motion was dynamically occluded. We further employed a subject-specific ROI-based multivariate pattern analysis investigating systematic BOLD modulations within low-level visual cortex associated with the spatial trajectory of occluded and visible targets. Focusing on V1, we tested whether changes in the V1-activity pattern are predictive of the stimulus trajectory during occlusion. In addition, we tested for similarities in informational content during visible and occluded stimulation, not only in V1, but also in neighbouring regions V2 and V3, plus within regions which have been related to motion processing (V5) and object identity (LO1, LO2) using probability maps of these regions. We hypothesised that a spatially-specific mental representation of different trajectories should result in an engagement of low-level visual areas, especially V1, during the occlusion phase. We also hypothesised that the activation pattern during the visible and dynamically occluded stimulus motion periods should be similar in low-level visual areas. Finally, we tested for the spatial layout in the modulated subregions of low-level visual areas using receptive field mapping. To anticipate, classification analyses yielded above chance accuracies already in the primary visual cortex when the classifier was trained on the visible data and tested on occluded data. The patterns of predictive informational content were highly similar to the pattern observed for visible trajectories and this result was further corroborated by visual field mapping.

## 2. MATERIAL & METHODS

### 2.1 Participants

Twenty-two right-handed participants (mean age=24.45, ±4.47, 14 women), with normal or corrected-to-normal vision, no history of psychiatric or neurological disorders and no regular intake of medications known to interact with central nervous system functions were recruited from the student community of Otto-von-Guericke Universität Magdeburg and gave informed consent to participate in the study, which was approved by the ethics committee of the Otto-von-Guericke-University. Participants could take part in the fMRI study only after successfully performing a velocity threshold determination task in a behavioural lab outside the scanner (see below for details). Six participants were excluded either due to poor performance during the main task (three with accuracy below 60%), absence on the last day of experiment (two participants) or poor quality of retinotopic mapping (one participant). The data of 16 participants (mean age=23.25, ±3.61, 12 women) were included in the final analysis.

### 2.2 General overview of the study design

The volunteers of this experiment performed a total of 8 tasks: a threshold determination task outside the scanner, one inside the scanner, twice familiarisation task with visible stimulation, twice prediction motion task with dynamically occluded stimulation, one independent functional localizer of the visual stimulation and a retinotopic mapping task. Here, we first present the chronological sequence of each task and below we describe the details of each paradigm. The first task was a behavioural threshold determination performed on a day prior to the scanning sessions, in which we ensured that participants could reliably perceive two different motion velocities. On day 1 of the scanning session, participants performed the same threshold determination task inside the scanner, to confirm the threshold of the previous session. This second threshold verification was followed by the functional localizer, and the first part of the main experiment (one session of familiarisation task followed by a session of the prediction motion task). On day 2, participants performed the 2nd part of the main experiment (again one session of familiarisation followed by prediction motion task), and the retinotopic and receptive field mapping.

### 2.3 Behavioural threshold determination: pre-scan session

Participants were placed in a dark room, 70 cm from the monitor (a 22-in., 120 Hz, LCD Screen, Samsung 2233RZ, recommended for vision research; Wang & Nikolić, 2011). All tasks were programmed using Psychophysics Toolbox (Version 3; Brainard, 1997) and run in Matlab 2012b (Mathworks Inc., Natick, MA, USA). On each trial, a white dot (1.24° visual angle) moved from the left to the centre (-6° to 0°) of a black screen. First, we presented a standard velocity of 16°/s (duration=300ms) which was randomly followed by one of the 11 possible velocities (t=t+t*0.05)^1^ including the standard velocity. The two moving stimuli were separated by an interstimulus interval (ISI) of 500 ms. Participants were instructed to keep their eyes fixed on the fixation cross (0.2°), attend to the movements and, after the disappearance of the second moving stimulus, to indicate whether the velocity of the second stimulus differed from the standard one. A total of 330 trials were presented divided in 6 blocks. The experiment lasted approximately 18 minutes. After the completion of the threshold experiment the data were fitted with a sigmoidal psychometric function (Curve Fitting Toolbox, Matlab, Mathworks Inc., Natick, MA, USA), and the time interval corresponding to 75% accuracy was estimated. This first task served to confirm that subjects were able to successfully discriminate velocities used in the main experiment. The same task was again performed inside the scanner in order to verify the participant’s threshold, which was used for all following prediction motion tasks inside the scanner.

### 2.4 Functional Localizer: Delineating low-level visual ROIs

The functional localizer was collected to identify areas in the visual cortex which responded to the visual stimulation used in our experimental runs and to later compare them with the location of the MVPA-spheres instrumental in motion extrapolation. We presented a high contrast checkerboard stimulus (1.6°) at seven different positions along the trajectories used in the main experiment (∼6.2°, ∼3.1°, ∼0°, horizontally and ∼6.2° and ∼3.1° vertically up- and downwards, while participants were asked to maintain fixation. The fixation cross (0.26°) was placed 14.2° to the right side of the stimulus central position. The stimulus flashed for 160ms, with an ISI that could vary from 1 to 2 seconds (Poisson-distributed) for 25 s at each position. Participants were asked to covertly attend the stimulation while keeping their eyes on the fixation cross. Fourteen blocks were separated in two runs, lasting in total approximately 7min20sec to complete the session.

### 2.5 Visible Phase: Association Learning during Visible Stimulation

During this initial association-learning phase, participants got familiarised with the main task by passively observing a stimulus moving first horizontally and then vertically on the screen (see Fig.1A). To this end, we used two velocities (fast = 300 ms, 20°/s or slow = individual participant’s threshold, on average 443.82 ms, ±24.20 ms or 14°/s) which were paired with two trajectories (upward or downward), leading to a 2 × 2 design with 4 possible combinations (order 1: up-fast, down-slow; order 2: down-fast, up-slow). The moving object consisted of a white dot (1.6°) moving from the left side of the screen to the centre (horizontal: 6.2° to 0°), then from the centre to the bottom (vertical: 0° to -6.2°) or top (vertical: 0° to +6.2°) of the screen. The stimulus was visible during the whole motion (in contrast to the occluded phase, see below). Participants completed a total of 100 trials divided into five runs, which in total lasted around 10 minutes. The ISI varied from 2 to 6 seconds (Poisson-distributed). No information about the velocity-direction association was provided to participants. The order of velocity-direction association was reversed on day 2 and the starting order was counterbalanced across subjects.

Participants were instructed to just observe the moving stimulus on the screen, with no further instruction for not priming them in any way. After the second run, we asked them, first, whether they had observed any regularities and, second, if they observed differences related to the direction-velocity information. We expected them to report the correct association (e.g. “when the dot moves fast it goes upward and when it moves slowly it goes downward”, or vice versa). If the participants didn’t report this relationship after the second run, they would be asked again^2^ after the next run. All participants completed a total of 5 runs of familiarisation.

### 2.6 Occluded Phase: Prediction of Dynamically Occluded Stimulation

Experimental set-up during testing was identical to the visible with the following exceptions: the moving stimulus was visible only during the horizontal movement and a grey rectangle (28.4° in width and 35.6° in height) was displayed during the whole run occluding the vertical trajectories. The respective end positions of the occluded vertical movements were marked by an “X” (1.6°) and the dot did not reappear. Participants performed a time and point of contact, i.e. prediction motion task. For this they were asked to respond when the moving stimulus would reach the top or bottom “X” mark (see Fig. 1B) and to indicate which of the two positions would be reached by pressing one of two buttons with their right index and middle finger. A total of 240 trials were presented divided in six runs. This experiment lasted approximately 33 minutes.

**Figure 1.**
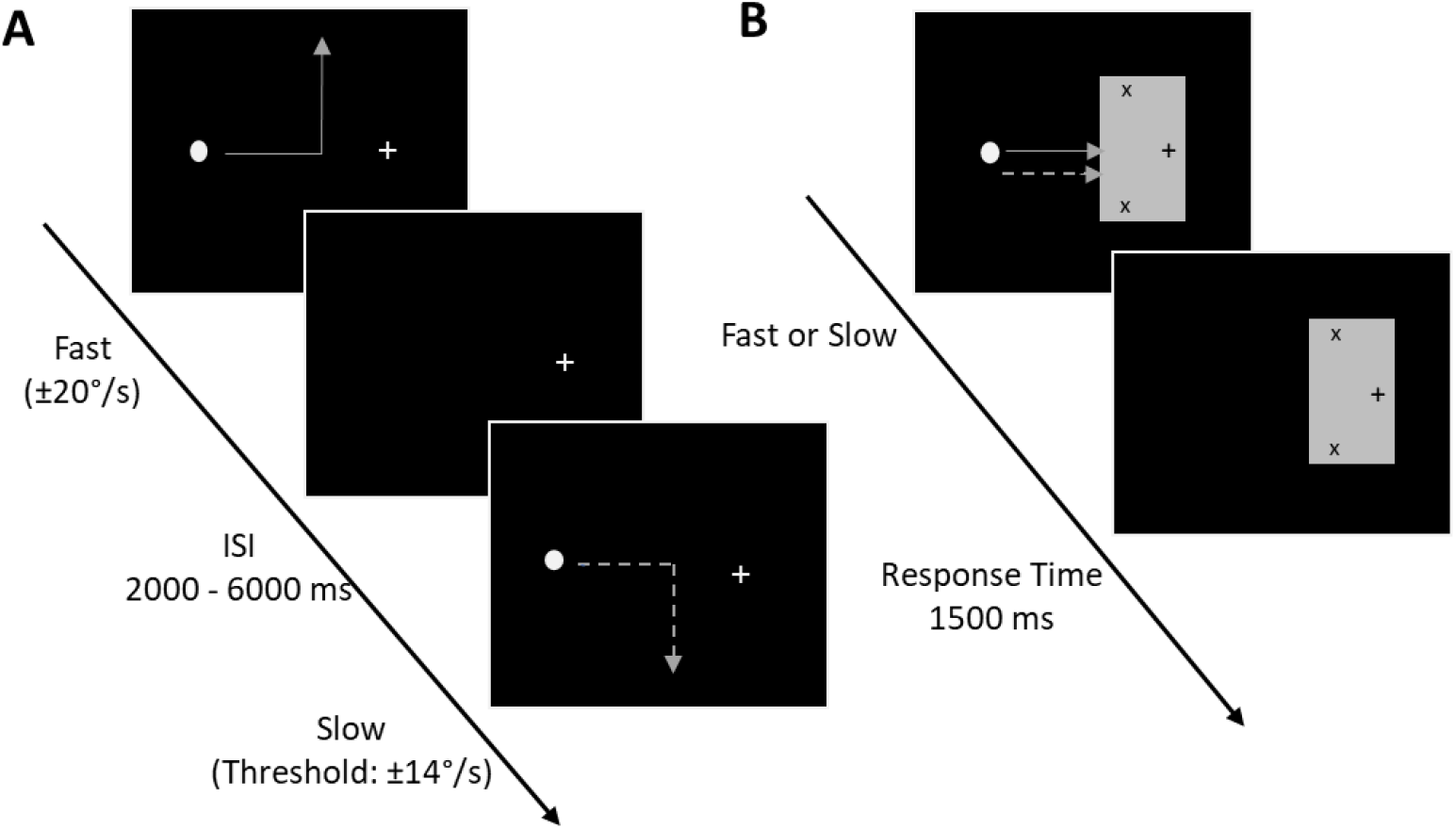
Display of the visual stimulation of the main experiment. (A) Visible Phase: Sequence of two trials observed by the participants. A white dot moved from the left side of the screen to the centre, then upwards or downwards as indicated by the arrows. The direction of the trajectory depended on the velocity of the dot here indicated by different types of line (solid and dashed). The full line represents fast movement and dashed line, slow movement. The lines are put here for illustrative purposes only, but were not displayed during the task. (B) Occluded Phase: The horizontal trajectories remained visible while vertical trajectories were occluded by a grey rectangle present during the whole trial. The “X” marks represented the stimulus final positions presented in the visible phase. Participants judged when and at which position the stimulus would end using the velocity information during the visible horizontal movement.

### 2.7 Retinotopic Mapping Phase

Seventeen participants were scanned in two sessions (nine participants performed the session on separate days due to maximal scanning time restrictions). The procedure used for measuring the retinotopic maps was adapted from Warnking et al. (2002) and Bordier et al. (2015). Stimuli were presented on a grey background. Visual eccentricity was mapped using a checkerboard ring which slowly contracted or expanded from the fixation dot. The speed of the expansion and the contraction varied linearly with the eccentricity (Bordier et al, 2015) and the ring reached a maximum diameter eccentricity of 6.6° and a minimum of 0.2°. When the maximum (expansion) or the minimum (contraction) was reached, a new ring would start from the original position. Polarity was mapped using one checkerboard wedge (10°) slowly rotating at a constant speed. Specific stimulation parameters were similar as the ones described by Warnking and colleagues (2002): the checkerboard stimulation flickered at a frequency of 8 Hz, in 10 cycles of 36 s each. The aspect ratio of the checkboards was kept constant (1.09) by scaling the height linearly with the eccentricity. In order to account for the effects of the hemodynamic signal, the wedges were presented clock- and counter-clockwise, and the rings were presented expanding annuli and contracting annuli (Warnking et al, 2002). In total, eight functional runs were acquired, two for each modality and direction, in two days, which in total lasted around 24 minutes per day.

## 3. APPARATUS & ACQUISITION

### 3.1 fMRI Experiment

The scanning sessions were conducted in a 3 Tesla Siemens PRISMA MR-system (Siemens, Erlangen, Germany), using a 64-channel head coil. The data of participants were acquired in 26 functional runs divided into two sessions, i.e. 210 volumes for the localization phase, 550 volumes for the visible phase and 1920 volumes for the occluded phase. Blood oxygenation level-dependent (BOLD) signals were acquired using a multi-band accelerated T2*-weighted echo-planar imaging (EPI) sequence (multi-band acceleration factor 2, repetition time (TR)=2000 ms, echo time (TE)=30 ms, flip angle=80°, field of view (FoV)=100 mm, voxel size=2.2 × 2.2 × 2.2 mm, no gap). Volumes were acquired in interleaved order. Identical slice selection on both days was achieved using the Head Scout Localizer which calculation is based on Autoalign (Siemens, Erlangen). Participants were placed inside the scanner and performed all tasks described above. Note that for threshold determination no fMRI data were collected. All visual stimuli were displayed on a rear-projection screen (302×170), approximately 350 mm from their eyes (± 10 mm depending on participant’s head size). Participants were asked to fixate on a cross (1.6°) and covertly attend to the stimuli. Fixation was controlled online during the whole fMRI-experiment using a fibre-optic camera (Kanowski et al., 2007).

In addition to the functional data a high-resolution three-dimensional T1-weighted anatomical map (TR = 2500 ms, TE = 2.82 ms, FoV = 256 mm, flip angle = 7°, voxel size = 1 × 1 × 1 mm, 192 slices, parallel imaging with a GRAPPA factor of 2, and 8 min scan duration) covering the whole brain was obtained using a magnetization-prepared rapid acquisition gradient echo (MPRAGE) sequence. This scan was used as a reference image to the EPI data during coregistration procedure and used as an overlay for the retinotopic and functional maps after inflation.

### 3.2 Retinotopy

Blood oxygenation level-dependent (BOLD) signals were acquired using a multi-band accelerated T2*-weighted EPI sequence (multi-band acceleration factor 2, TR=2000 ms, TE=30 ms, flip angle=90°, FoV=128 mm, voxel size=2.2 × 2.2 × 2.2 mm, no gap). For each run, 180 volumes were acquired in interleaved order.

A high-resolution three-dimensional T1-weighted anatomical map was obtained only for the occipital lobe (TR=2500 ms, TE=2.82 ms, FoV=256 mm, flip angle=7°, voxel size=1 × 1 × 1 mm, 192 slices, parallel imaging with a GRAPPA factor of 2, and 8 min scan duration) using a magnetization-prepared rapid acquisition gradient echo (MPRAGE) sequence. This scan was used as anatomical reference to the EPI data during the registration procedure.

## 4. STATISTICAL ANALYSIS

### 4.1 Behaviour

Subjects’ temporal and spatial estimates of target stimulation were measured by subjects’ response time after stimulus occlusion and correct prediction of vertical direction (accuracy), respectively. Missed trials, trials with RT smaller than 0.1 s and greater than the mean + 3 std were excluded from the analysis. In addition, we calculated the temporal estimation error by taking the difference between the amount of time the stimulus travels behind the occluder and participants’ response time. We used a 2×2 repeated measures ANOVA (direction × velocity) for investigating differences in accuracy, response time and temporal estimation error. All analyses were calculated using JASP (v. 0.15.0, https://jasp-stats.org/). JASP was also used to compute post hoc tests (simple main effects function) and effect size (partial η2).

### 4.2 Retinotopy

A three-dimensional reconstruction of the cortical sheet based on the structural image of each of the 16 subjects was performed using the recon-all function of Freesurfer (https://surfer.nmr.mgh.harvard.edu/). Retinotopic maps along the polar and eccentricity dimensions were calculated for each of the cortical surfaces using the “selxavg3-sess” function of Freesurfer. The right-hemispheric low-level visual areas V1, V2, V3 were delineated manually on the flattened cortical sheets based on the boundaries of phase reversals within the polar angle maps (Abdollahi et al. 2014). Delineation of borders were created based on of Georgieva et al. (2009) and Kolster et al. (2010). Based on these delineations, we created six masks of V1, V2 and V3 for upper and lower visual areas, which later was used to identify region-specific local maxima during the visible and occluded phase. Probabilistic maps of MT and LOs as provided by Freesurfer parcellation for each subject were included in the analyses. Freesurfer labels were converted to volume ROIs based on Freesurfer mri_vol2roi function. LO masks were separated in LO1 and LO2 using Georgieva et al. (2009) and Kolster et al. (2010) delimitations. Results of the LO and MT ROIs can be found in the supplementary material.

### 4.3 fMRI preprocessing

Participants’ data from both days were analysed using SPM12 (www.fil.ion.ucl.ac.uk/spm, Wellcome Trust Centre for Neuroimaging, London, UK). The first five volumes of each run were discarded to allow for steady state magnetization. Functional images were slice-timing corrected and spatially realigned (registered to the mean image). Head motion parameters were later used as nuisance regressors in the general linear model (GLM). Finally, the structural image was coregistered (estimate and reslice) to first functional image of the first run. Resliced images were smoothed with a gaussian kernel of 6 mm.

### 4.4 fMRI data Modelling

The participants’ functional data of day 1 and day 2 for each task were modelled with a single general linear model (GLM, Friston et al, 1995), which included the run-wise condition parameters, derivatives, and six motion regressors as nuisance covariates. In particular, regressors of each condition (up-fast, down-slow, up-slow, down-fast or up-slow, down-fast, up-fast, down-slow) were specified by using canonical hemodynamic response function (HRF). Temporal and dispersion derivatives of each regressor were added to the model in order to account for variability in the onset response (Friston et al., 1998). From the condition-specific maps of beta weights averaged across runs of each participant, we extracted beta weights from subject-specific V1, V2, V3 (see below for details of retinotopic analysis) for the univariate group analysis using MarsBar 0.44 (Brett, et al., 2002). A 2×2×2×3 repeated measures ANOVA was calculated with the factors: direction (upward, downward), velocity (fast/slow), visual region quadrants (VQ; upper/lower) and visual regions (V1, V2, V3) for both visible and occluded phases, followed by post-hoc analyses, when necessary.

### 4.5 Multivariate Pattern Analysis

In order to complement the univariate analysis, we executed a series of multivariate pattern analyses, using CoSMoMVPA (Oosterhof, Connolly, and Haxby, 2016), designed to identify whether patterns of activity during visible stimulation can be used to accurately classify the patterns of activity during occlusion. To this end, we performed volume-based searchlight analyses with a 2 mm radius using run-wise beta weights (proportional to percent signal change), in native space for each condition, as datasets (two beta-values per run: one for each condition). Searchlight analyses were chosen to retain a high spatial specificity. In particular, a linear discriminant analysis (LDA) classifier was trained in the 10 runs of the visible phase (20 beta values), using a leave-one-run-out approach, and tested in the 12 runs of the occluded phase (24 beta values). As sanity checks, the classifier was also trained and tested on the runs of visible only and occluded only, using n fold partitions. As a further manipulation check, the classifier was also trained and tested on motion velocity (see supplementary material for this last sanity check).

We carried out one searchlight analysis per region of interest (upper and lower V1-V3 etc.), to increase the spatial specificity to the classification and to be able to draw conclusions per functional region. For each searchlight within a given region one accuracy value was obtained. We focused the analysis on the specific spheres inside the ROIs which contained informative voxels and, for that, we adopted a thresholding procedure. We applied a cut-off allowing only values above 0.5 to be in the analysis, excluding chance level spheres, as we predicted that only a restricted number of searchlight spheres would contain meaningful information, and computed the average of the 5% highest accuracy values of the distribution. Our choice of selection criterion is also in agreement with a recent publication, suggesting that consciously perceived stimulus information is not solely restricted to the regions within visual cortex directly coding the stimulus (van Kemenade, et al., 2021). Hence, by using an information-based rather than a visual-localizer based criterion, we ensured to have a spatially unbiased selection criterion (see below next section 4.6. for an independent assessment of this novel thresholding approach introduced here).

To evaluate the statistical significance for each ROIs, permutation tests was carried out for each subject. The permutation included 1,000 iterations which contained randomised data labels per run, keeping the same original dataset. For a spatially accurate comparison, we obtained the accuracy value from the same searchlight spheres included in the 5% highest accuracy sample, for all 1000 samples for each individual person. The 5% maximum values were averaged across spheres for the original and permuted dataset permutation. For group level analysis, we followed the Etzel (2017) approach. The null distribution contained the average across participants for each of the 1000 permutations with the addition of the true-labelled group-level average, resulting in 1001 group-level accuracies. The permutation p value was computed by taking the sum of the permuted accuracies higher or equal to the true-labelled accuracy and dividing by the number of iterations plus 1.

### 4.6 Projecting the spatial layout of MVPA-results on visual field maps

By utilizing the well-established knowledge that low-level visual cortex is spatially organised, we tested whether the searchlight spheres with the highest decoding accuracy were overlapping across the independent statistical tests (decoding of visual motion, occluded motion and occluded motion by visual motion). The rationale behind that was to confirm whether the regions inside each visual area, which encoded trajectory information, were common across visible and occluded phases. To this end, statistical maps were first projected onto the flattened cortical sheets. Based on the specific polar and eccentricity maps, each vertex of the univariate and multivariate result maps could be associated with a specific location within the visual field and later be overlaid for comparison. Hence, in addition to showing flattened anatomical maps, visual field maps were used as well for a standardized projection of the results independent on individual cortical sheets. We restricted this analysis to lower and upper V1, as these regions contain neurons with the smallest receptive field size (Barbot, Xue & Carrasco, 2021). An overlap of significantly modulated spheres would suggest at least spatial proximity during the processing of the visible stimuli and during visual extrapolation. To further corroborate these findings, the univariate results of the functional localizer phase (contrast: up vs down condition) were also overlaid, to confirm independently that the regions containing patterns of significant MVPA-results were in close proximity to the areas responding to visual stimulation in the independent functional localizer.

## 5. RESULTS

### 5.1 Behavioural read-outs

#### Spatial Estimation (Where)

After the visible phase, we expected the participants to accurately indicate the motion direction according to the velocity-direction association. Results corroborate our expectations, indicating that participants had very high-performance accuracy (Figure 2. A). Group averages for all occluded conditions were above .92 accuracy (up-fast: mean (M)=.951, ±.035; up-slow: M=.969, ±.025; down-fast: M=.923, ± .070; down-slow: M=.952, ±.039). Main effects of direction and velocity were not significant (F(1,15)=3.734, p=0.072, ɳ2 =.199; F(1,15)=3.851, p=0.069, ɳ2=.204, respectively). No interaction between factors (F(1,15)=.318, p=.581, ɳ2 = .021) was observed. These non-significant differences across conditions observed here may suggest that the association learning had the same level of difficulty independently of direction and velocity.

**Figure 2.**
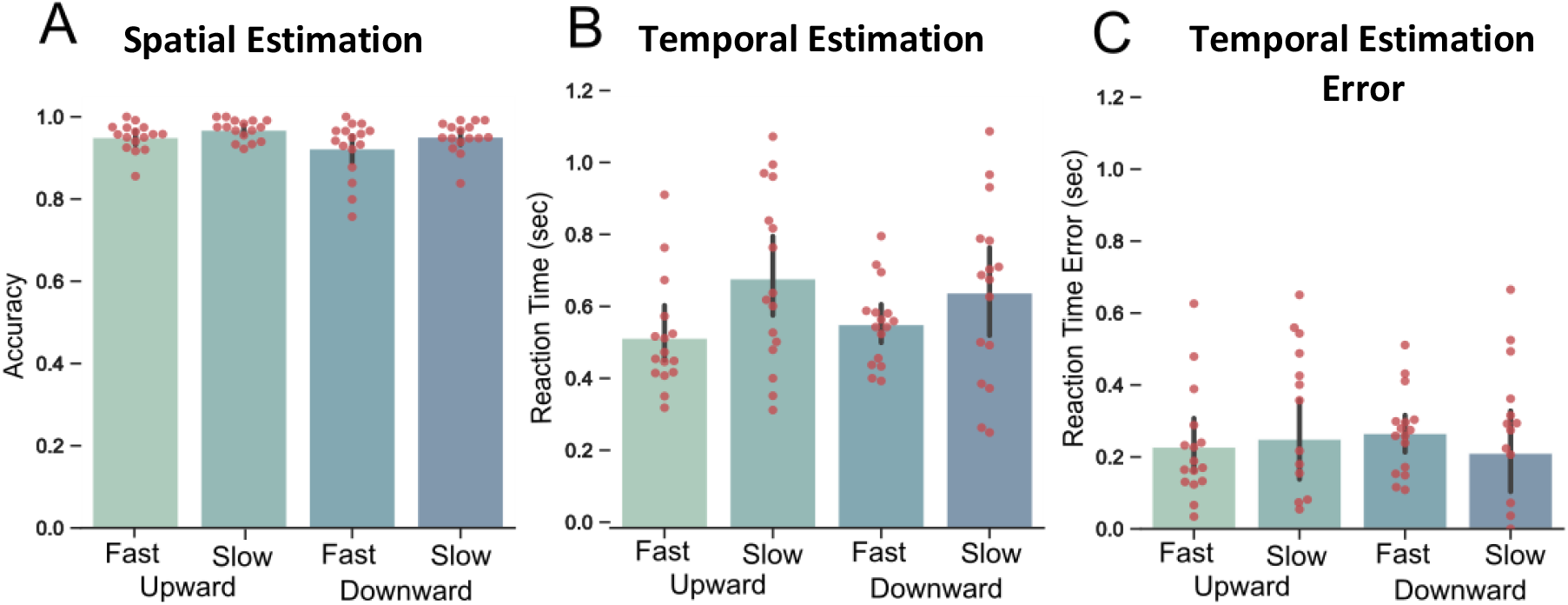
Behavioural results: In all bar graphs (from left to right), light green bar (1^st^ bar) depicts fast condition, darker green (2^nd^ bar) slow condition in upward direction, light blue bar (3^rd^ bar) depicts fast condition and dark blue bar (4^th^ bar) slow condition in downward direction. Red dots superimposed on each bar represent behavioural results of all individual subjects. (A) Group average accuracy for spatial estimation. (B) Group average reaction times for temporal estimations. (C) Group average reaction times for temporal estimation error (difference between physical stimulus displacement time and estimated time).

#### Temporal Estimation (When)

Participants showed a high consistency in their time estimation (Figure 2.B): Average response times (RT) were consistent with physical stimulus velocity, i.e. the time-to-contact in the slow condition was estimated to be later than the fast condition (up-fast: M=.528, ±.154; up-slow: M=.694, ±.244; down-fast: M=.566, ±.114; down-slow: M=.654, ±.247). Accordingly, a main effect of velocity (F(1,15)=8.704, p=.010, ɳ^2^=.367) was observed. No significant main effect was found for direction (F(1,15)=.003, p=.957, ɳ^2^=.000), as well as no interaction between factors (F(1,15)=1.059, p=.320, ɳ^2^=.066).

#### Temporal Estimation Error

Figure 2.C shows similar estimation errors across all conditions (up-fast: M=.228, ±.154; up-slow: M=.250, ±.241; down-fast: M=.266, ±.114; down-slow: M=.211, ± .243). In accord, no significant main effect for direction (F(1,15)=.003, p=.957, ɳ^2^=.000) or velocity (F(1,15)=.157, p=.698, ɳ^2^=.010), and also no interaction (F(1,15)=1.059, p=.320, ɳ^2^=.066) was observed.

The results above indicate that, at the behavioural level, participants accurately estimated the stimulus end position, as well as time-to-contact, the latter with a certain yet consistent bias from the reality across all conditions (temporal overestimation).

### 5.2 Univariate activation mapping

#### 5.2.1 Subject-specific results

In the first analysis step, we identified modulations in fMRI signal for each individual participant using univariate analysis. Comparisons of the trajectories (upward vs. downward) revealed significant patterns of activity in regions representing upper and lower visual quadrants contralateral to the stimulated hemifield, as expected (see Fig. 3 for an exemplary subject; all other subjects can be found in the supplementary Figure 1). In contrast, the comparison of speed levels (fast vs. slow) did not yield any significant modulations of fMRI signals in low-level visual areas.

**Figure 3.**
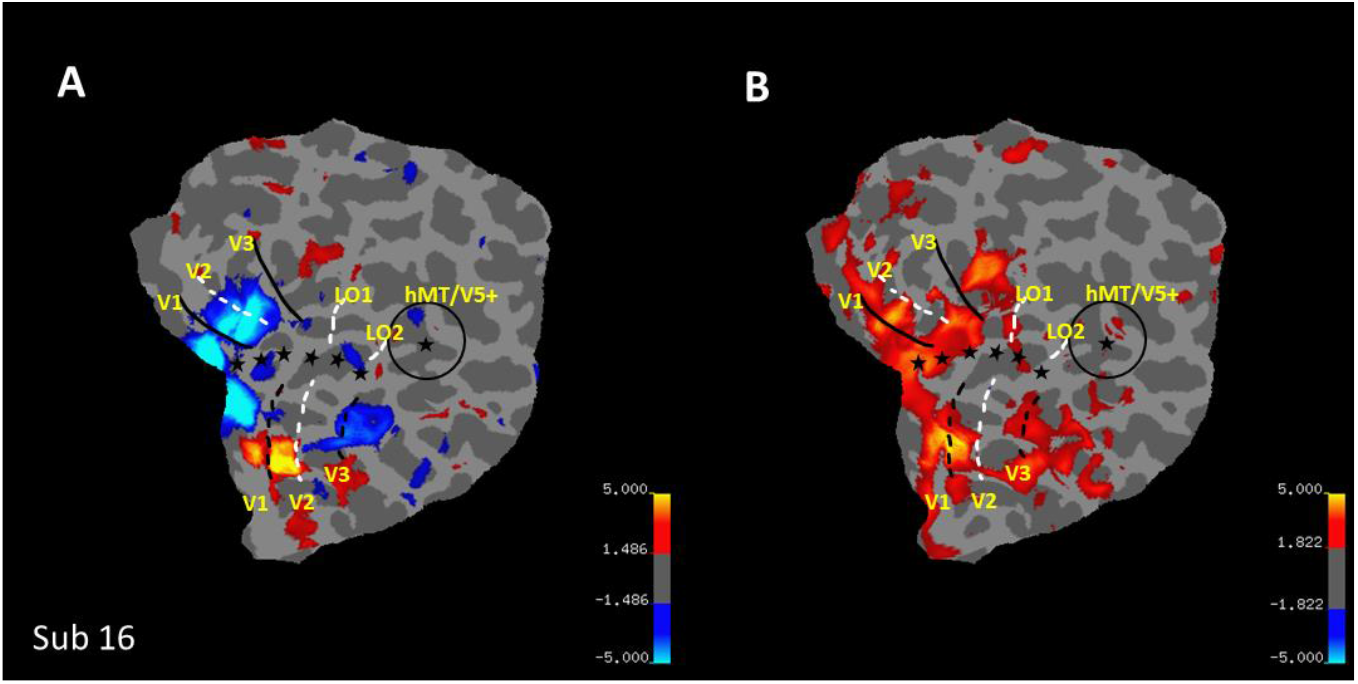
Univariate results of an exemplary participant during (A) Visible Phase and (B) Occluded Phase, for contrast between upward (warm colours) vs. downward (cold colours) projected on the individual flat map. Retinotopic map delimitations are indicated by stars (central visual field), plus white and black full and dashed lines indicating borders between visual fields (Abdollahi et al., 2014).

#### 5.2.2 Group results

##### Statistical comparison of univariate Modulations during Visible & Occluded Phases

From all low-level regions of each individual participant based on retinotopic masks, we extracted beta weights during the visible phase from the local maxima and averaged them across participants (Fig. 4A). To test – in a first manipulation check – if we would be able to differentiate upward and downward trajectories of visible stimuli in V1-V3, we then statistically compared these beta weights. During the presence of the target moving upward in the visible phase, *lower V1, V2 and V3* showed enhanced fMRI-signals compared to downward movements (lower V1-upward: M=8.376, ±3.845 vs. downward: M=2.985, ±2.818; V2-upward: M=7.081, ±3.880 vs. downward: M=1.868, ±2.092; V3-upward: M=6.450, ±3.165 vs. downward: M=1.998, ±2.010), as shown by the paired one-tail Student’s T-tests (upward > downward: t(15)=7.534, p_holm_<.001; t(15)=6.655, p_holm_<.001; t(15)=5.998, p_holm_<.001, respectively). During downward movements in *upper V1* compared to upward movements the analysis revealed similar results (upper V1-downward: M=11.797, ±5.845 vs. upward: M=3.773, ±3.738; V2-downward: M=9.418, ±4.229 vs. upward: M=1.749, ±2.251; V3-downward: M=7.788, ±3.162 vs. upward: M=1.381, ±2.457), (V1-; V2-; V3-),, as revealed by the paired one-tail Student’s T-tests (downward > upward: t(15)=7.944, p_holm_<.001; t(15)=6.902, p_holm_<.001; t(15)=8.653, p_holm_<.001, respectively). Together, these findings show that the visual stimulation was salient enough to elicit differential fMRI-signals in accord with motion direction.

**Figure 4.**
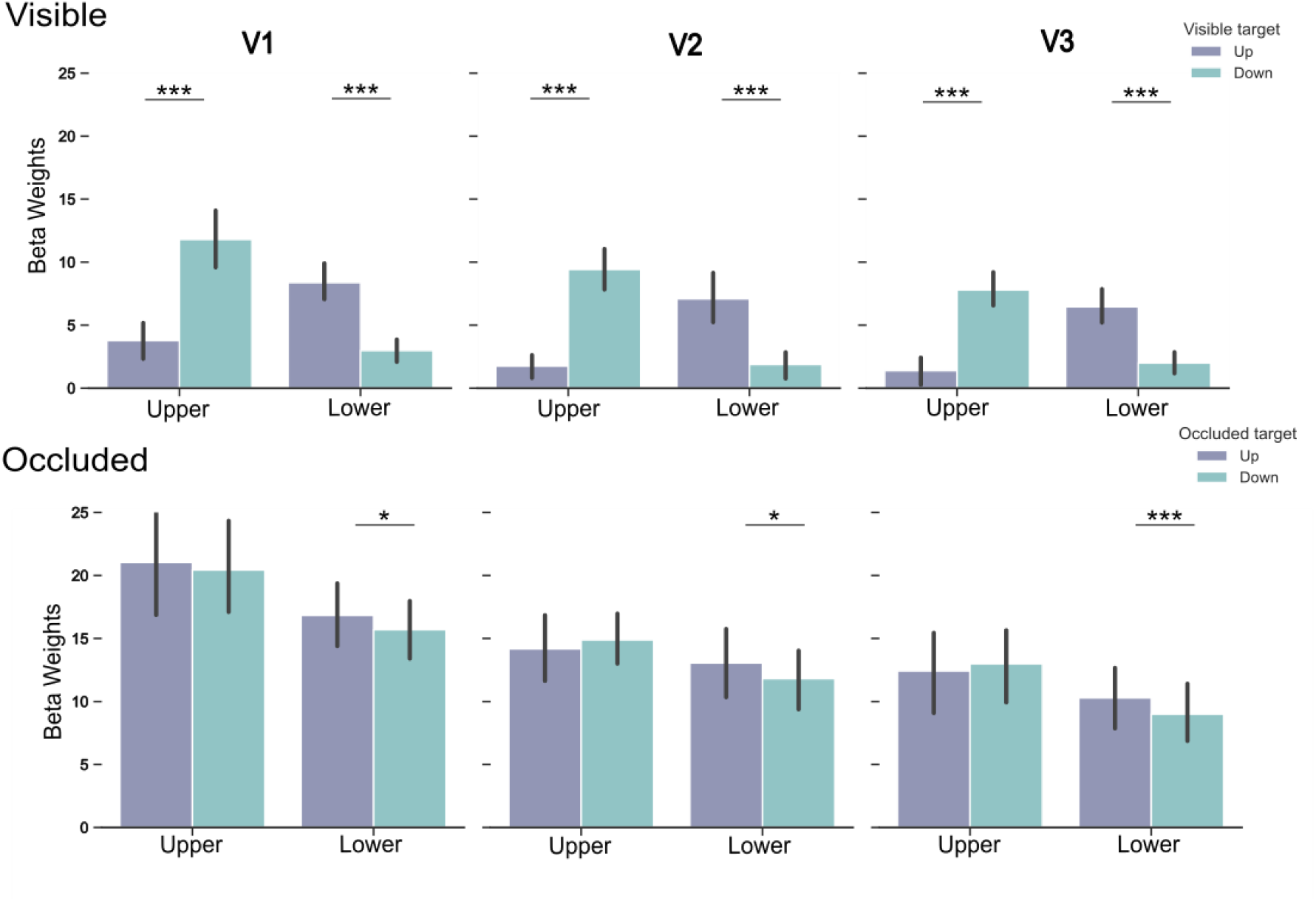
Univariate beta weights (proportional to percent signal change) during visible phase (upper row) and occluded phase (lower row). Green bars depict average beta weights for downward trajectories, while purple bars average beta weights for upward trajectories. Stars indicate significance between conditions inside each region on interest.

The identical locations independently identified during visible stimulation (see above) were then used to analyse activation patterns during occlusion. During the occluded phase, a pattern similar to the visible phase was present for the lower VF (Figure 4B): When participants were asked to temporally and spatially estimate upward movements, enhanced fMRI-signals compared to downward movements were observed in *lower V1* (lower V1-upward: M=16.834, ±6.225 vs. downward-: M=15.696 ±6.428; V2-upward: M=13.057, ±7.234 vs. downward: M=11.806, ± 6.725; V3-upward: M=10.283, ±6.777 vs. downward: M=8.997, ±6.381). Statistical analyses revealed significant differences in all three regions (upward > downward: t(15)=2.261; p_holm_=.040; t(15)=1.990, p_holm_=.040; t(15)=2.919, p_holm_=.015, respectively). Unexpectedly, higher response during upward movements was also registered in the *upper V1* (upper V1-upward: M=21.027, ±11.401 vs. downward: M=20.435, ±10.059); however, this difference was not significant (downward > upwards: t(15)=-0.851; p=.796). In contrast, we found the hypothesised pattern for upper V2 (upper V2-upward M=14.174, ±5.896 vs. downward: M=14.891, ±5.061) and V3-upward: M=12.426, ±8.558 vs. downward: M=12.986, ±8.072). Yet, these results were not significantly different (downward > upwards – upper V2: t(15)=1.023, p=.161; upper V3: t(15)=-0.677, p=.254).

##### Unfolding interaction between factors

While the t-tests above were pre-planned to directly test our main hypothesis, below we extend our analysis by including the factor velocity, visual area (V1, V2, V3) and visual quadrant (upper VQ = upper V1-V3; lower VQ = lower V1-V3) using repeated measures ANOVAs. In the visible phase statistical analysis revealed significant interaction between direction and VQ (F(1,15)=131.862, p<.001, ɳ^2^=.898), plus velocity and direction (F(1,15)=5.582, p=.032, ɳ^2^=.271) and velocity and VQ (F(1,15)=30.566, p<.001, ɳ^2^=.671). Post-hoc tests confirmed that fMRI-responses, due to downward direction, were higher than upward direction in the upper VQ (MD= -7.367, SE=.672, t=-10.962, p_bonf_<.001) and responses due to upward directions were higher in the lower VQ compared to downward direction responses in these regions (MD=5.019, SE=.672, t=7.468, p_bonf_ <.001), confirming the above t-tests. In addition, slow motion resulted in significantly higher fMRI-signals in upper relative to lower visual quadrant; while no such effect was seen for fast motion (VQ *velocity - slow in upper VQ vs. slow in lower VQ: MD=5.940, SE=0.589, t=4.991, p_bonf_ <.001) and the slow motion also differed for upward vs. downward direction, while no such effect was observed for fast motion (direction*velocity: up-slow vs. down-slow: MD=-2.185, SE=.586, t= -3.726, p_bonf_ =.005). Finally, the distinct pattern of responses yielded main effects for direction (F(1,15) = 8.575, p=.010, ɳ^2^=.364), VQ (F(1,15)=5.744, p=.030, ɳ^2^=.277, and visual areas (F(1,15)=9.327, p<.001, ɳ^2^=.383) with higher responses for the downward vs. upward direction, higher responses for upper VF field than lower VF and enhanced responses in V1 compared to V2 (MD=1.704, SE=.558, t=3.053, p_bonf_=.014) and to V3 (MD = 2.328, SE = .558, t=4.172, p_bonf_ =<.001).

During the occlusion phase, statistical analysis again revealed interactions between direction and VQ (F(1,15)=5.331, p=.036, ɳ^2^=.262), supporting the t-test results, direction and velocity (F(1,15)=32.746, p<.001, ɳ^2^=.686), velocity and visual areas (F(1,15)=5.371, p=.010, ɳ^2^=.264), plus a triple interaction between direction, velocity and visual areas (F(1.22,18.4)=19.087, p<.001 (Greenhouse-Geisser-corrected), ɳ^2^=.560). A Post-hoc comparison for the hypothesized interaction indicated marginally significant results for downward motion in the lower visual quadrant compared to upper visual quadrant (MD=3.938, SE=1.412, t=2.788, p_bonf_ =.077). In addition, post-hoc tests for interaction of direction and velocity revealed that fast motion led to enhanced responses compared to slow motion in the upward direction (MD=5.129, SE=1.280, t=4.006, p_bonf_=.003), while no significant effect for fast vs. slow motion was observed for the downward direction. Post-hoc tests of velocity*visual areas revealed that fast responses compared to slow response were most prominent in V1, whereas the triple interaction with direction revealed that this elevation in V1 was highest for upward motion, while significant differences for downward motion was seen only in V1 compared to V3 for fast condition (see supplementary Figure 2. Finally, main effects revealed results similar to the visible phase with upper visual regions eliciting higher responses than lower visual regions (F(1,15)=5.439, p=.034, ɳ^2^ = .266). Moreover, V1, V2 and V3 again expressed differential effects (F(1,15)=16.534, p<.001, ɳ^2^=.524). Comparisons showed that V1 presented higher beta values than V2 (MD=5.016, SE=1.302, t=3.851, p_bonf_ =.002) and V3 (MD=7.325, SE=1.302, t=5.624, p_bonf_ <.001).

Together, the univariate results indicated that low-level visual regions were modulated by direction at least in one of the VQ during occlusion phase, thus they do in part support our hypothesis. However, the absence of significant effects in one quadrant might be due to the lower sensitivity of univariate analysis approaches as it is well known that multivariate pattern analysis has a higher sensitivity than traditional univariate analysis (e.g. Li et al., 2009). Moreover, MVPA allows for drawing conclusions about the representational content within activation patterns (Anzellotti & Coutanche, 2018). To test our research question even more thoroughly, we performed run-wise GLM analysis and used the resulting beta values as input to multivariate pattern analysis.

### 5.3 Multivariate Pattern Analysis

A series of volume-based MVPA analyses was applied to test for pattern similarity between visible and occluded stimulation periods.

#### Classifying Direction Patterns of Visible from Occluded Phase

This classification analysis was most crucial to test our hypothesis for a common engagement of low-level visual cortex during the presentation of visible and dynamically occluded motion. It was performed with the main purpose of decoding similar activation patterns for visible and occluded phases in upper and lower V1-3. For these multivariate analyses, maps of beta weights for upward and downward motion were calculated run-wise separately for visible and occluded phase and used as input.

Results (Fig.5 and table 1.A) show that direction-specific informational patterns from the visible phase could be used to decode informational patterns in the occluded phase in the lower AND upper V1, V2 and V3 significantly extending the results from the univariate analysis by showing the similarity in spatial layout of the informational content of these two phases (see below for a more thorough description of the spatial layout within V1).

**Figure 5.**
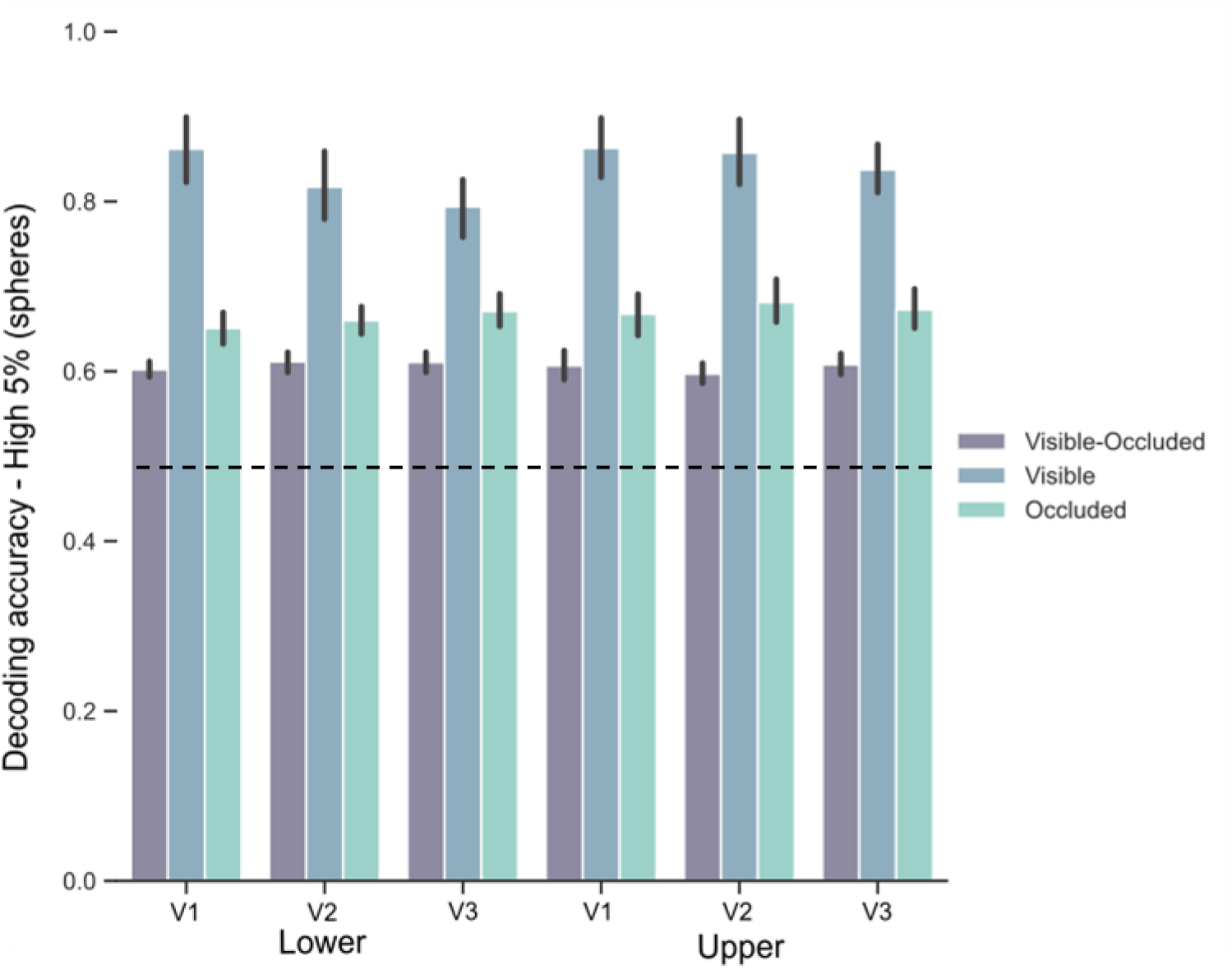
Decoding accuracies for all classification analyses in upper and lower V1-V3. Dashed line depicts the theoretical chance level, though note that the chance level used for statistical testing was derived from permutations tests. Purple bars show average accuracies for the classification analysis trained on visible data and tested on occluded data. Blue bars show the accuracies for the classification analysis with training and testing the visible phase only and green bars depict average accuracies for the analysis using occluded data also during both training and testing.

**Table 1.**
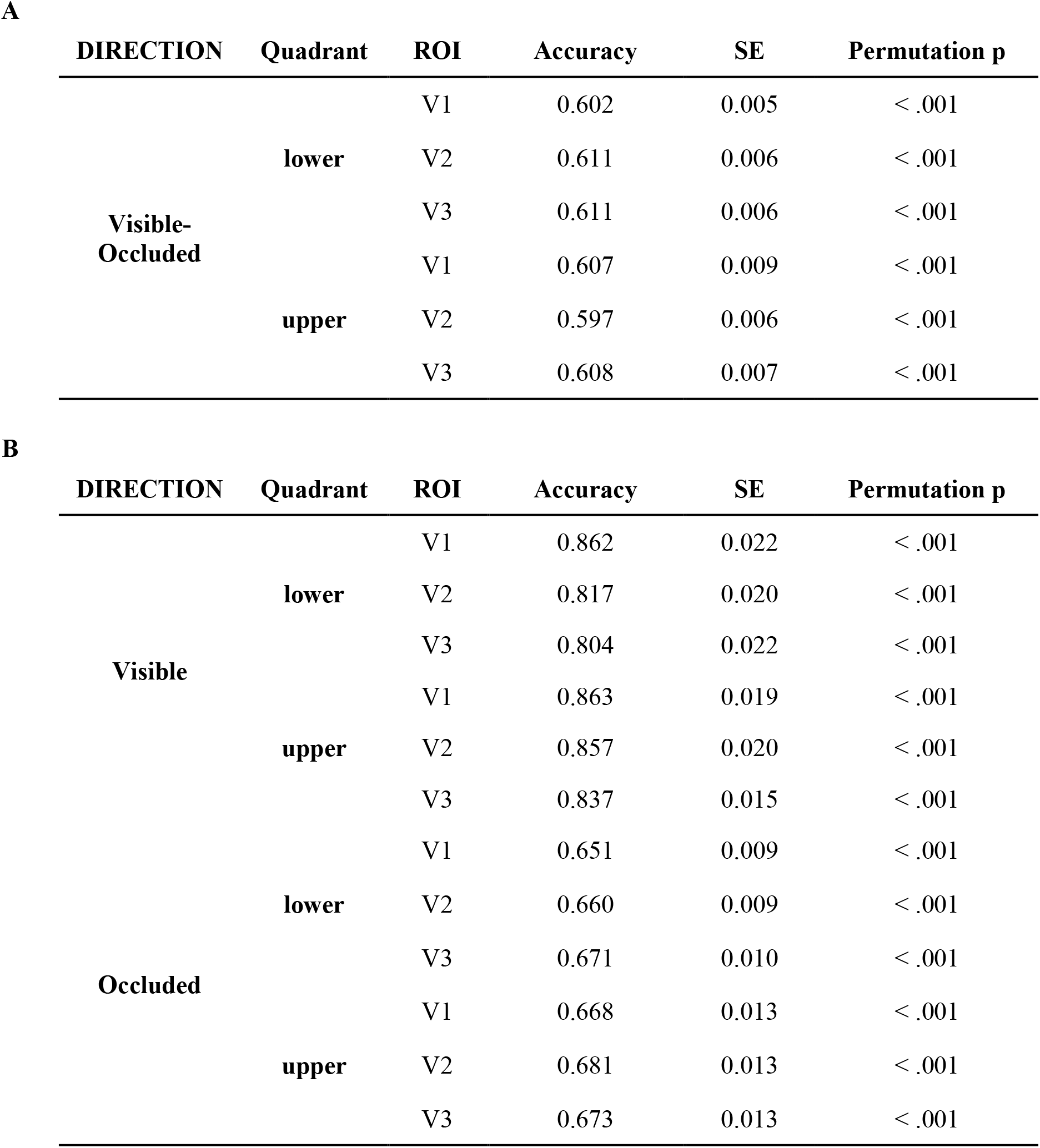
Decoding accuracies for (A) visible-occluded classification analysis and (B) separate visible and occluded classification analyses, in upper and lower V1-V3.

#### Classifying Direction Patterns from Visible & Occluded Phase

As manipulation checks, we performed MVPAs separately for the visible and occluded phase to check for reliability of the classification. For both phases we found direction-specific informational patterns of activity predicting motion trajectory. For these two separate analyses, we also expected accuracy values to be higher when the classifier was trained and tested in the visible phase, once the stimulus was all the time present. Indeed, results (Table 1.B) indicated higher accuracy values for visible phase compared to the cross-phase analysis and to the occluded phase analysis, and higher values of the latter compared to the cross-phase analysis (Fig.5). These results further confirmed that the classifier decoded the relevant direction-specific information and that the information was somewhat diluted during occlusion compared to visible stimulation.

Above, we used averaged scores of accuracies across participants. For maximal transparency and to give the reader an impression of the interindividual variability, Figure 6 shows the two relevant measures average decoding accuracies (top) and number of spheres for each ROI (from which average decoding accuracies are calculated, Fig. 6 bottom), for every subject. We observed a higher interindividual variability in decoding accuracy during the occluded phase, compared to the other phases. The number of spheres included in the 5% sample of most informative voxels adopted also varied across subjects, following a heterogeneous pattern, but did not show a bias towards one analysis. The average of spheres across the three analyses is 84.21 (±30.08) spheres.

**Figure 6.**
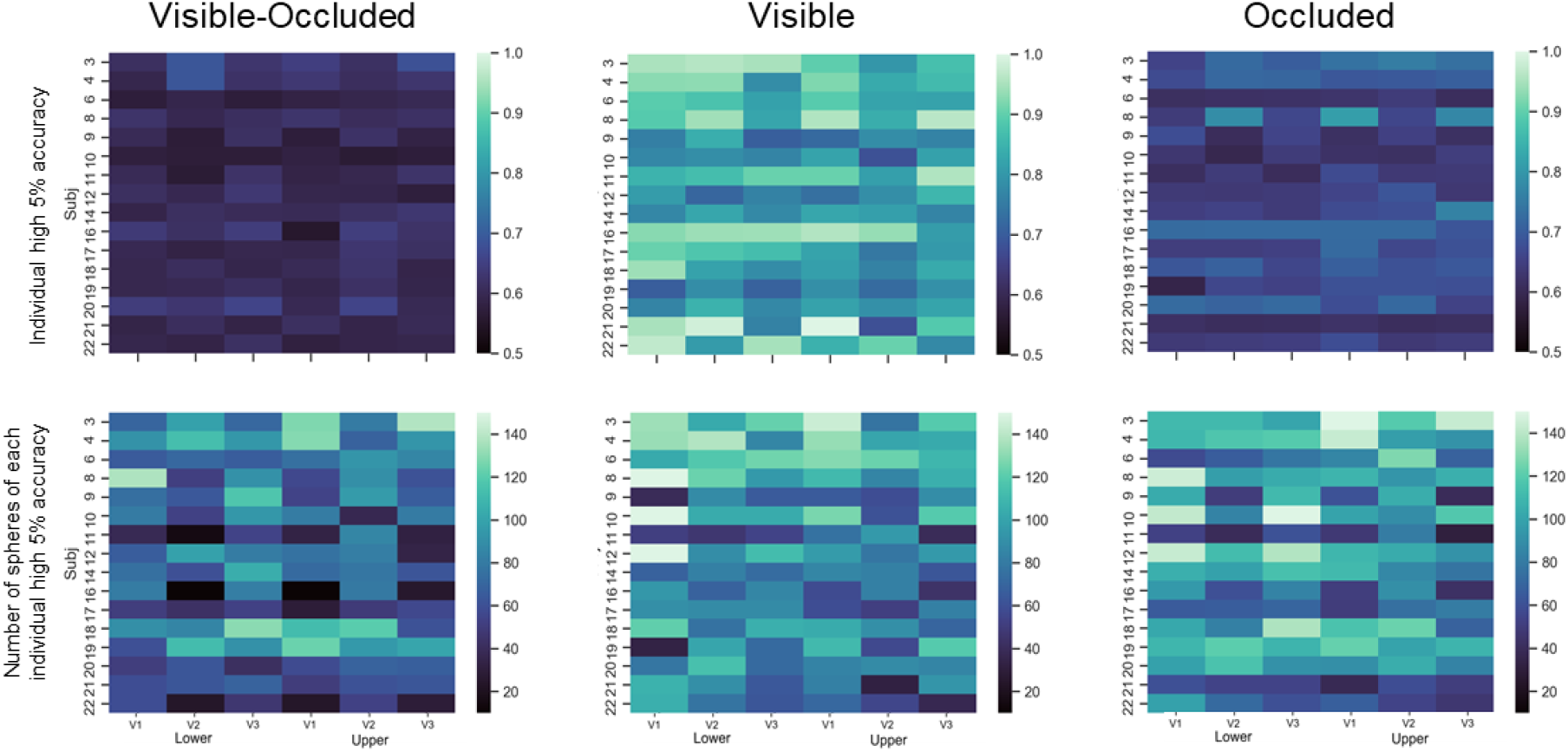
Upper row depicts average decoding accuracies of each participant in each of the 6 ROIs. The lower row depicts the number of spheres included in the calculation of the 5% highest accuracy values, for each subject in each ROI.

#### Spatial layout of significant spheres across classification analyses

The projection of location of the significant spheres map for each analysis onto the retinotopic maps was carried out to verify their exact anatomical location. Comparing the overlap of spheres maps across analyses will tell us where, within the visual field, the map can be found and what their exact spatial layout would be, i.e. scattered or concentrated. This, in turn, allows us to draw firm conclusions regarding their representational content, especially if they are concentrated in a portion within the visual field which encodes the visible stimulus. For this, we focused in V1 which contains neurons with the highest spatial acuity and projected retinotopic maps on visual field representations (Duncan & Boynton, 2003; Song et. al, 2015). Moreover, we also included the results from the univariate functional localiser as a further manipulation check. For the visual fields subject-specific density maps were computed signifying the concentration of significant spheres within specific regions of the visual field. Figure 7 (top) shows visual field maps of an exemplary subject, in which we observed overlapping density maps for the different statistical tests. Across participants, overlap of the functional localiser (green line) with significant spheres from the visible phase (blue line) was observed for 13 participants in lower V1 and for 8 participants in upper V1, which indicates a robust reliability, once the stimulus is physically present even across multivariate and univariate methods. For visible and occluded phases, the overlap was observed for all 16 in both upper and lower V1, while for visible and visible-occluded, we observed the overlap for 14 participants in the upper V1 and 15 for the lower V1. Finally, for occluded and visible-occluded phases, the overlap was found for 13 in the upper V1 and 14 participants in lower V1. It should be noted, however, that for some subjects we find that parts of the density map fall outside the stimulated visual quadrant. This scattering was most likely caused by the quality of the polarity maps which was not high enough in all parts of visual cortex to allow for an errorless transformation, though note that the delimitation of different visual fields was not affected by these local variations in polar maps. Importantly, when inspecting the location of significant spheres in anatomical space overlaid onto the curvature, polarity and eccentricity maps (see Fig. 7-bottom), we could confirm that the significant clusters were located in the upper and lower lip of V1, respectively.

**Figure 7.**
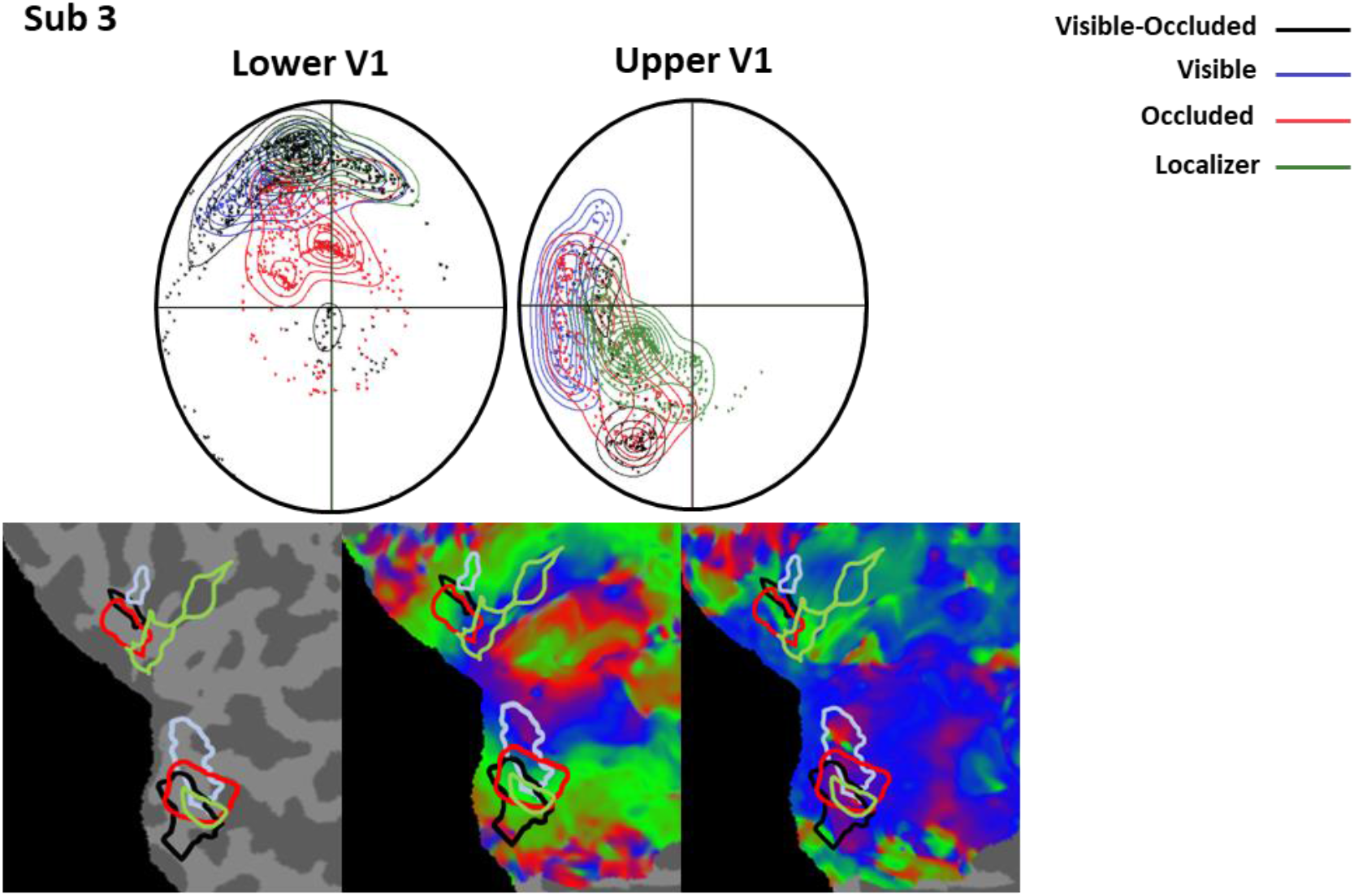
Significant decoding accuracy maps projected onto retinotopic maps and derived visual field maps of one exemplary subject. Top: Visual fields of lower V1 and upper V1. Black curvatures and dots depict the localization of the vertices with significant accuracies for the visible-occluded classification. Blue and red curvatures and dots represent the localization of the vertices with significant accuracies for the visible and occluded classifications, respectively. Green lines and dots represent the same, but here results from the univariate functional localizer was used. Dots which spread to other quadrants could indicate scattered representations, but might be partly reflect the quality of retinotopic mapping itself. Bottom: Significant decoding accuracy maps overlaid on flattened anatomical maps of occipital cortex (right), the retinotopic eccentricity (middle) and polarity maps (left).

We interpret our findings as evidence that different types of information can be found in similar regions of the visual field within primary visual cortex. Results for different analyses of spheres, selective for different types of visual stimulation, converge within topographically organized regions within the early visual cortex. Importantly, it should be noted that this pattern of results is unbiased as the different analyses were spatially unrestricted within visual regions and carried out with independent data sets from different runs, at least for the analysis of the functional localiser, the visual only and occluded only MVPAs. That confirms that the information-based searchlight selection criterion did not randomly pick arbitrary voxels within the visual cortex. Rather, the selection reflected the spatial selectivity of cortical regions representing the very same stimulus throughout different conditions, even during occlusion.

## 6. DISCUSSION

Our study tested the involvement of low-level visual regions in continuous motion extrapolation by comparing visible and occluded stimulus trajectories. Our behavioural results demonstrate that participants were highly accurate in judging the target’s direction and time of arrival. The general overestimation of time intervals that we observed seems to be related to our specific stimulus settings. Here, short intervals (around 250-800 ms) were employed which are often overestimated (Benguigui, Ripoll, Broderick, 2003; Bennet, Uji, Baurès, 2018; Vicovaro, Noventa, Battaglini, 2019). Remarkably, univariate fMRI-results showed that, to some extent, activity in low-level visual regions was tied to estimated motion trajectories. The more sensitive MVPA-analysis revealed that the activity pattern in V1 evoked by visible motion was indeed informative about the direction of the trajectory during occlusion. Finally, the location of informative spheres was consistent across several independent classifications, strongly suggesting that the sub-regions within low-level visual areas, coding the stimulus trajectory, were selectively engaged; and this claim was further supported using fMRI-based receptive field mapping of the informative regions.

### 6.1. Univariate fMRI-results

Our *univariate* fMRI-analysis compared velocity and direction in low-level visual areas. Results revealed that velocity did not differentially modulate fMRI-signals in any low-level areas, while the motion trajectory selectively modulated neural activity during the visible phase and partially during occlusion. We attribute this outcome to the dominance of spatial estimation over temporal estimation, which might be due to the well-known higher spatial acuity of the visual system (Klein et al, 2018, Welch & Warren, 1986). Our motion direction results indicate that low-level visual areas were engaged during the occlusion of the target, thereby extending previous observations on the involvement of primary visual cortex in apparent motion (Ekman, Kok and Lang, 2017) to continuous motion extrapolation. In particular, the vertical visible trajectory elicited responses in the upper visual area for downward direction and in the lower visual area for upward direction, as expected. During the dynamic occlusion of the stimulus, similar patterns of activation in the lower quadrants of the visual regions were observed, partly supporting particular theories about the mechanisms of motion extrapolation, which posit that a visual representation of a non-visible object is maintained during occlusion. These results are also in line with previous mental imagery findings which investigated imagery using retinotopic mapping and found activation in both striate and extrastriate cortex for imagined objects (Slotnick, et al, 2005).

However, this similarity in univariate activation patterns was less pronounced for the upper visual areas (lower visual field), rather an overall higher activation in upper V1 relative to lower V1 was observed regardless of visual trajectory. Asymmetry in the upper vs. lower visual quadrants has been previously reported and one possible explanation might be the distribution of ‘near-preferring’ neurons, which tend to be more frequent in lower VF compared to upper VF (Nasr & Tootell, 2018, 2020; Karim & Kojima, 2017). In general, psychophysiological studies demonstrated that lower visual field seems to be more thoroughly engaged in tasks from different domains, such as motion (Danckert & Goodale, 2001; Levine & McAnany, 2005, Lakha & Humphreys, 2005), colour discrimination and hue sensitivity (Levine & McAnany, 2005). This difference between lower and upper visual field is known as vertical meridian asymmetry which is more pronounced at larger eccentricities (Carrasco, Talgar & Cameron, 2001, Barbot, Xue & Carrasco, 2021). Visual field asymmetries have also been reported for the crowding phenomenon (He, Cavanagh, & Intriligator, 1996), the spatial resolution of attention (Intriligator & Cavanagh, 2001), distribution of receptive field properties and orientation preference (Merkel, 2020) and may be further amplified by reading habits (Rinaldi, Di Luca, Henik, & Girelli, 2014). Finally, there might have been an evolutionary advantage preferring downward over upward movements, i.e. for catching things falling down than flying away, similar to the preference for looming vs. receding stimuli (e.g. Tyll et al., 2011), which could explain the asymmetry observed for the univariate results.

In addition to the upper vs. lower VF asymmetry observed during the comparison between visible and occluded vertical trajectories, we also observed overall enhanced fMRI-signals during partially occluded relative to the visible phase. Note that participants passively observed the stimulus travelling on the screen during the visible phase, whereas during the occluded phase, they actively had to be engaged with the task. This latter task set may have required a higher level of attention (Zuanazzi & Noppeney, 2020; Klein, Harvey, Dumoulin, 2014). On the theoretical level, this difference may be explained in the light of the Rao and Ballard (1999) predictive coding model. In the visible phase, once the stimulus became predictable, less error-correction signals might have been exchanged between lower and higher visual areas, reducing the height of the neural response (Alink et al, 2010). On the other hand, the lack of visual information during the occluded phase might have intrinsically decreased the predictability level, leading to higher activity due to the need of prediction. On a mechanistic level, the activity in V1 during occlusion may have also been primarily caused by feedback projections from higher cortical regions terminating in supramarginal layers, whereas during passive viewing feedforward projections in the laminar layer may have been primarily engaged. Accordingly, fMRI-responses in supramarginal layers was recently found to be enhanced relative to lower cortical layers (de Hollander et al.,2020).

In contrast to our univariate results, previous fMRI studies often failed to reliably observe the recruitment of low-level visual areas during motion prediction (Schuwaiti, Curtis & Johnson (2007) Olson and colleagues (2003) observed high engagement of inferior parietal sulcus and also reported a strong decrease in fMRI-signal in V1/V2 during occlusion. Our contradictory results could be attributed to the use of techniques such as retinotopy, which allowed us to precisely identify low-level visual areas individually in each participant, targeting the specific low-level visual regions, while accounting for inter-individual anatomical variability (Greenlee, 2000, Amunts et al. 2000). In accord with our results, two recent studies which based their analysis on subject-specific retinotopic mapping were able to observe effects in low-level visual cortex during imagery of motion (Emmerling et al, 2016) and shape predictions (Erlikhman & Caplovitz, 2017). Both studies also used multivariate pattern analysis, which allows to draw conclusions not only on correlative brain-behaviour relationships but on the informational content represented within brain regions.

### 6.2 Multivariate fMRI-results

Our MVPA results further extend our findings by showing that the patterns of activity in upper vs. lower visual field representations could be used to predict encoded motion trajectories during visible stimulation and during occlusion on their own. Most importantly, the data from visual trajectory could also be used for training to predict invisible motion during occlusion. It should be noted that these results significantly extend the univariate results during the occlusion phase by showing that the pattern of activity in low-level visual areas shows some commonalities during visible motion and occlusion. These results are in line with a recent study which suggests that even with no stimulation in these regions, it is possible to decode information based on continuous perception (van Kemenade et al, 2022). Additionally, the successful prediction does not depend on the difference in activation height for upward vs downward tasks, as the lower visual regions did not show enhanced fMRI-signals for occluded upward vs. downward motion in the univariate analysis. Therefore, the pattern of activity, rather than an overall difference in response-amplitude most likely accounts for the results in V1 and also in V5 (see supplemental material), which did not show significant univariate results (see also Wang et al., 2014 for similar results), but yielded significant classification accuracies for motion direction and velocity during occlusion using MVPA.

Despite the similarities in activation pattern, which led to the significant prediction accuracies, it should also be noted that there were some differences in MVPA-results between occluded and visible conditions: group averaged accuracy values were systematically higher for visible classification than occluded classification. This difference may attribute to the fact that the classifier was capturing a more reliable response with visible stimulation compared to a dynamically stimulus occlusion.

Remarkably, the projection of significant spheres onto the retinotopic maps suggested that the location of the most informative voxels overlapped across the three independent classification analyses (visible and occluded stimuli, plus across category) plus the univariate functional localizer. This overlap of visible and occluded informative activity patterns in the visual field may suggest that shared computational circuits in primary visual cortex do exist, supporting both processes. However, layer-specific fMRI and/or single-cell studies are needed to further corroborate this interpretation. The spatial layout of our results also confirms the plausibility of our sphere-selection approach, since overlapping results were found across different analyses. Our subject-specific approach was necessary once the target regions show considerable interindividual anatomical variability, thus differing from the standard fMRI-group analysis, which is traditionally based on the assumption of spatial commonalities across participants after spatial transformation of individual brains to a standard reference brain.

Finally, a remarkable outcome from this study is that our results were similar to mental imagery studies (e.g. Albers et al., 2013), despite the fact that no explicit instruction was given to the participants to imagine the stimulation. Instead, we aimed to investigate whether low-level visual cortex would engage during the presentation of occluded stimulation without the participant having previous knowledge about a specific strategy for performing the task. Future studies could explicitly instruct participants to imagine occluded trajectories and observe if the observed effects vary as a function of task instruction.

To conclude, we investigated the neurobiological processes underlying motion extrapolation as indexed by patterns of activity in low-level visual regions during the presentation of visible and dynamically occluded trajectories. Our data support the notion that virtually identical regions inside lower and upper V1, V2 and V3 represent information about visual stimulus trajectories in the absence of visual stimulation and suggests that shared neural circuits may be utilized when processing visible and extrapolated trajectories.

## Acknowledgement

We would like to thank for the support we received from Denise Scheermann and Claus Tempelmann at the fMRI lab from the Department of Neurology at Otto-von-Guericke Universität-Magdeburg.

## CRediT author statement

**Camila Agostino:**Conceptualization, Software, Formal Analysis, Investigation, Writing-Original Draft, Visualization. **Christian Merkel**: Software, Formal Analysis, Writing - Original Draft. **Felix Ball, Peter Vavra:** Writing – Review and Editing. **Herman Hinrichs:** Resources, Writing – Review and Editing, Funding acquisition. **Toemme Noesselt**: Conceptualization, Supervision, Writing - Original Draft, Funding acquisition.

## Footnotes

1 The intervals used for slow stimulation were taken from the set of milliseconds: in degrees/second: {16, 15.23, 14.51, 13.8, 13.16,12.53, 11.93, 11.37, 10.82, 10.31, 9.822}.

2 All participants were able to report the association after the third run. Some didn’t understand exactly what they had to report after the second run. However, after observing the third run, they all reported the correct association.

