## Supplementary Material for "Seeing and Extrapolating motion trajectories share common informative activation patterns in primary visual cortex"

### 1. Univariate Analysis

#### 1.1. Subject-specific results

Figure 1 depicts modulations in fMRI signals during Visible (left maps) and Occluded (right maps) phases. Trajectory comparisons (upward vs. downward) revealed increased patter of activity in regions representing upper and lower visual quadrants opposite to the stimulated visual field, as expected.

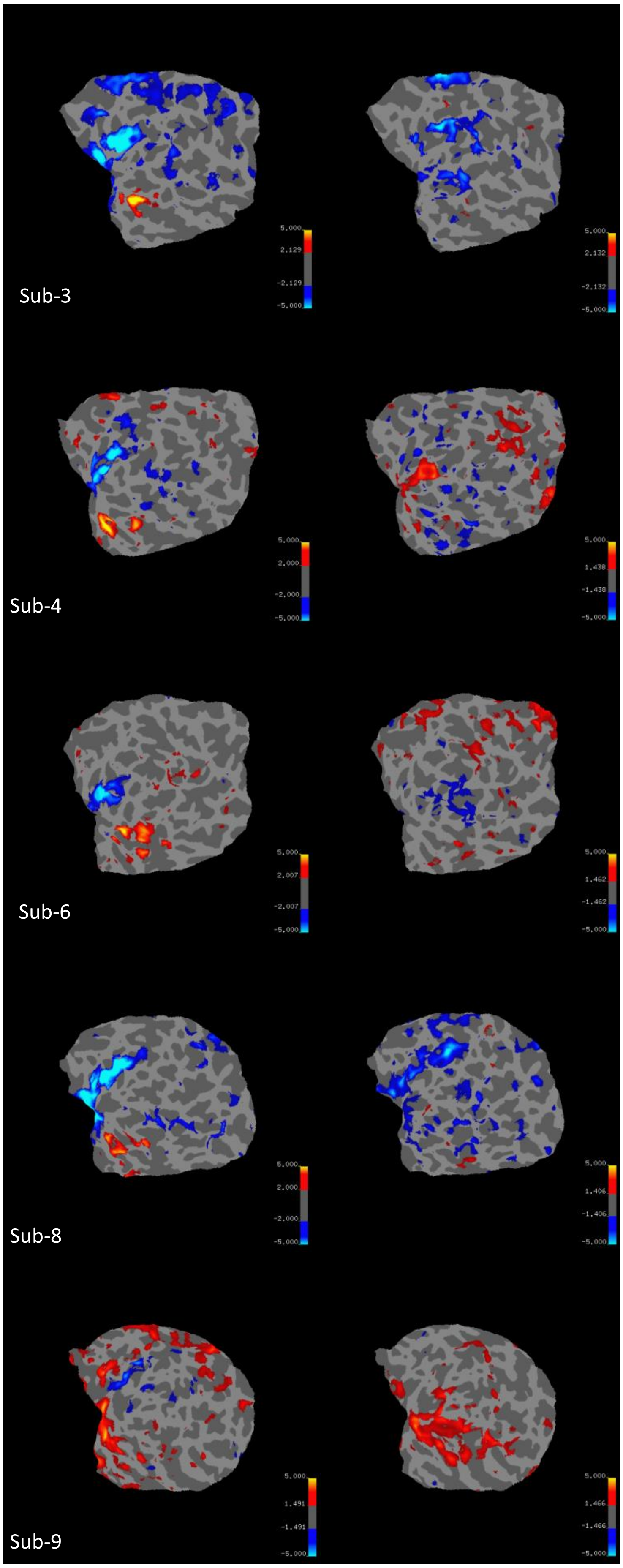
Sub-3
5.000
2.129
-2.129
-5.000
5.000
2.132
-2.132
-5.000
Sub-4
5.000
2.000
-2.000
-5.000
5.000
1.438
-1.438
-5.000
Sub-6
5.000
2.007
-2.007
-5.000
5.000
1.462
-1.462
-5.000
Sub-8
5.000
2.000
-2.000
-5.000
5.000
1.406
-1.406
-5.000
Sub-9
5.000
1.491
-1.491
-5.000
5.000
1.466
-1.466
-5.000

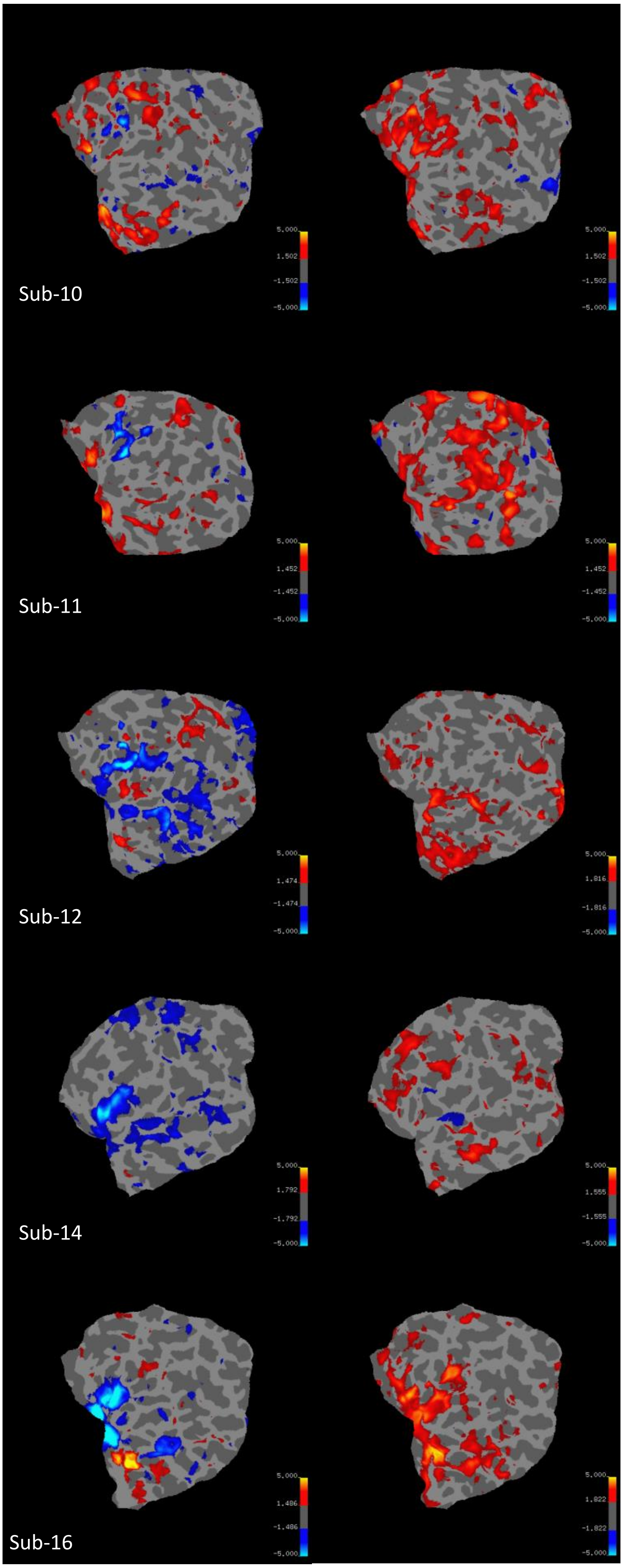
Sub-10
5.000
1.502
-1.502
-5.000
Sub-11
5.000
1.452
-1.452
-5.000
Sub-12
5.000
1.474
-1.474
-5.000
1.816
-1.816
Sub-14
5.000
1.792
-1.792
-5.000
1.555
-1.555
Sub-16
5.000
1.486
-1.486
-5.000
1.822
-1.822

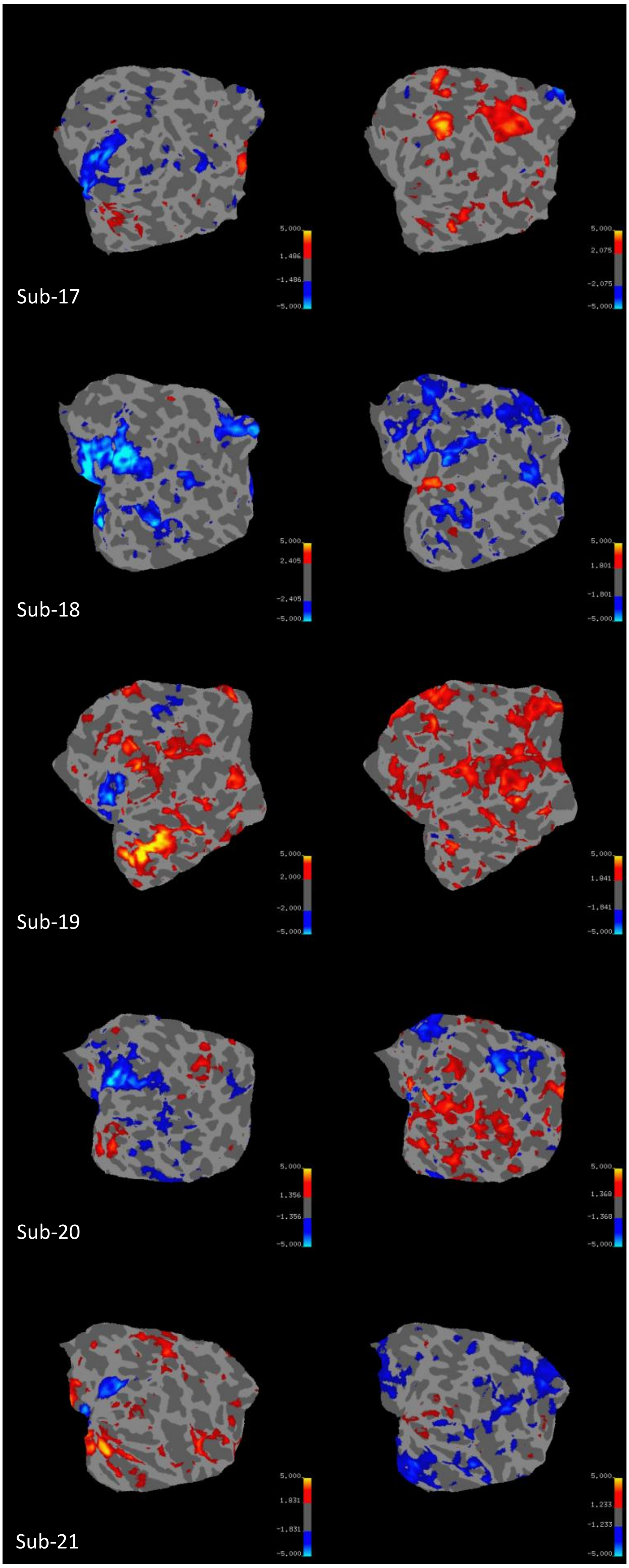

5,000
1,486
-1,486
-5,000
Sub-17
5,000
2,075
-2,075
-5,000
5,000
2,405
-2,405
-5,000
Sub-18
5,000
1,801
-1,801
-5,000
5,000
2,000
-2,000
-5,000
Sub-19
5,000
1,841
-1,841
-5,000
5,000
1,356
-1,356
-5,000
Sub-20
5,000
1,368
-1,368
-5,000
5,000
1,831
-1,831
-5,000
Sub-21
5,000
1,233
-1,233
-5,000

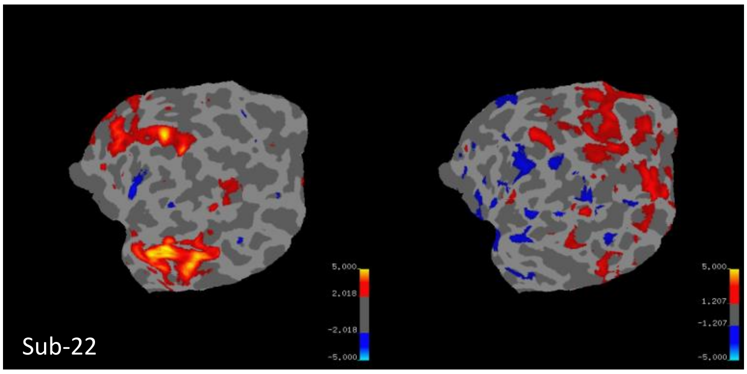


*Suppl. Figure 1 - Univariate results of all participants, during (A) Visible Phase and (B) Occluded Phase, for the contrast upward (warm colours) vs. downward (cold colours) motion projected on the individual flat map. Visual flat maps were created by cutting V1 along the fundus of the calcarine figure. However, this fundus was not the border between lower and upper field representations in all subjects (see, for instance subject 10, whose upper visual field representations reach into the upper lip of the calcarine sulcus). Therefore, activations corresponding to upper and lower field representations are not always restricted to the upper and lower anatomical part of low-level visual areas, but appear to spread into the other quadrant as well.*

#### 1.2. Group-specific results

Post-hoc tests indicated that significant differences from the interaction between velocity and visual fields (VFs) resulted from fast motion in V1 compared to V2 (MD=6.349, SE=1.386, t=4.581, $p_{bonf}<.001$) and V1 compared to V3 (MD=8.681, SE=1.386, t=6.264, $p_{bonf}<.001$). Comparison from the triple interaction (velocity*VFs*direction) showed significant differences, mainly in V1 (see suppl. Fig. 2). Fast motion in upward direction was more prominent compared to fast motion in downward direction (MD=5.912, SE=0.837, t=7.061, $p_{bonf}<.001$) and to slow motion in upward direction (MD=8.549, SE=1.419, t=6.025, $p_{bonf}<.001$). Fast motion in upward direction also modulated higher fMRI-signals in V1 compared to V2 (MD=7.665, SE=1.418, t=5.405, $p_{bonf}<.001$) and V3 (MD=10.356, SE=1.418, t=7.303, $p_{bonf}<.001$). In contrast, differences in fast motion was also observed in downward direction in V1 compared to V3 (MD=7.006, SE=1.418, t=4.940, $p_{bonf}<.001$).

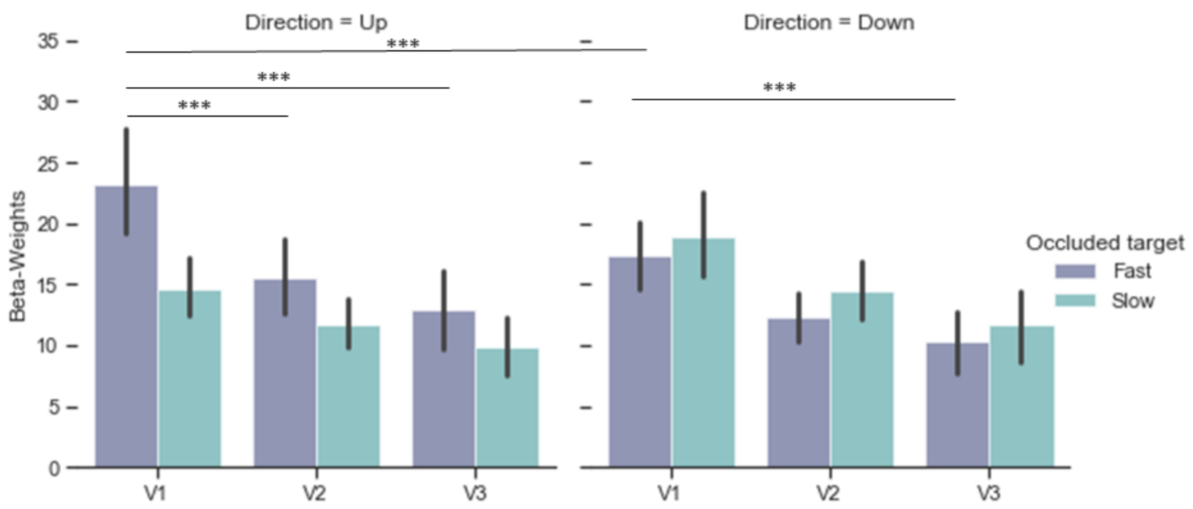


*Suppl. Figure 2 - Univariate beta weights (proportional to percent signal change) occluded phase. Purple bars (from left to right) depict average beta weights for fast motion, while green bars, the average beta weights for slow motion. Stars indicate significance between conditions inside each region on interest*

**2. Multivariate Pattern Analysis - *Complementary Analysis***

In addition to the low-level visual ROIs V1-V3, we also performed classification analyses with LO1, LO2 and hMT/V5+, for easy comparison with previous results.

*Classifying additional ROIs:* We trained a classifier on the distinction of up vs. downward condition during the visible phase and tested on the occluded up- vs. downward trajectories. Accuracy levels above chance were found in right LO1 (Accuracy (Acc)=0.589, SE=0.007, permutation p ($p_{perm}$)<.001), right LO2(Acc=0.597, SE=0.007, $p_{perm}$ <.001), right hMT/V5+ (Acc=0.597, SE=0.006, $p_{perm}$<.001) and left (Acc=0.588, SE=0.005, $p_{perm}$<.001). Results suggest that these regions also carried a similar pattern of information found in low-level visual areas.

*Classifying Velocity Patterns of Visible from Occluded Phase*: Here we carried out a classification analysis training in visible and testing in occluded phases (same procedure used in the first described analysis), but setting fast and slow as targets. Results indicated accuracies significantly above chance in lower V1 (Acc=0.607, SE=0.005, $p_{perm}$<.001), upper V1 (Acc=0.597, SE=0.006, $p_{perm}$<.001) and bilateral hMT/V5 (rh: Acc=0.608, SE=0.004, $p_{perm}$<.001, lh: Acc=0.605, SE=0.006, $p_{perm}$<.001).
